# Domestication and Temperature Modulate Gene Expression Signatures and Growth in the Australasian Snapper *Chrysophrys Auratus*

**DOI:** 10.1101/387084

**Authors:** Maren Wellenreuther, Jérémy Le Luyer, Denham Cook, Peter A. Ritchie, Louis Bernatchez

## Abstract

**ABSTRACT**

Identifying genes and pathways involved in domestication is critical to understand how species change in response to human-induced selection pressures. We experimentally manipulated temperature conditions for F_1_-hatchery and wild Australasian snapper (*Chrysophrys auratus*) for 18 days and measured differences in growth, white muscle RNA transcription and haematological blood parameters. Over 2.2 Gb paired-end reads were assembled *de novo* for a total set of 33,017 transcripts (N50 = 2,804). We found pronounced growth and gene expression differences between wild and domesticated individuals related to global developmental and immune pathways. Temperature modulated growth responses were linked to major pathways affecting metabolism, cell regulation and signalling. This study is the first step towards gaining an understanding of the changes occurring in the early stages of domestication, and the mechanisms underlying thermal adaptation and associated growth in poikilothermic vertebrates. Our study further provides the first transcriptome resources for studying biological questions in this non-model fish species.

## INTRODUCTION

The domestication of plants and animals marks a major evolutionary transition and ascertaining the molecular and physiological basis of domestication and breeding represents a vibrant area of interdisciplinary research (Zeder 2015). Compared to terrestrial animals, the domestication of fish for human consumption started only recently (Yáñez *et al.* 2015) and with the exception of a few species, such as the common carp (*Cyprinus carpio*) or Nile tilapia (*Oreochromis niloticus*), most domestication effort date back to the early 1980s (Balon 2004). Consequently, most cultured fish species have only slightly changed from their wild conspecifics (Olesen *et al.* 2003; Li and Ponzoni 2015). This represents a unique opportunity to study how animals evolve during the transition from wild to captive conditions, as well as during the first generations of domestication.

For pokilothermic species such as fish, temperature plays a profound and controlling role in their biological functioning (Fry 1971; Hochachka and Somero 2002). Affecting cellular components via a change in the viscosity of body fluids, fluidity of cell membranes, and enzyme kinetics (Currie and Schulte 2014) temperature influences the pathways by which individuals allocate energy to competing functions (Claireaux and Lefrançois 2007; Khan *et al.* 2015). For eurythermal fish (which can survive across a broad temperature range), such as the Australasian snapper (*Chrysophrys auratus*, Sparidae), environmental fluctuations dictate that their body temperatures vary in both space and time. When environmental temperatures are within a ‘zone of tolerance’ physiological, biochemical and behavioural aspects of the organisms biology are at, or near, optimal (Pörtner 2010; Currie and Schulte 2014). Yet when temperatures are at the extremities of this tolerable range both acute and chronic stress responses can be observed and translate into reduced organismal performance, adversely affecting growth, routine activity, or reproduction (Fry 1971; Mininni *et al.* 2014; Schulte 2014). However, the thermal tolerances of fish commonly show significant plasticity, with notable intraspecific variability and acclimatory responses reported in both eurythermal and stenothermal (narrow thermal tolerance) species (Seebacher *et al.* 2005; Anttila *et al.* 2013; Sandblom *et al.* 2014; Metzger and Schulte 2018). Given the profound influence of temperature on fish metabolism and organismal performance, a comparison of how temperature affects wild and domestic strains of snapper is an important question to address.

Rapid growth is a key determinant of commercial farming success, and is heavily modulated by the ambient temperature (Mininni *et al.* 2014; Bizuayehu *et al.* 2015; Besson *et al.* 2016). Moreover, growth is frequently correlated with a number of life-history traits, such as gonad maturation and reproductive timing (Schaffer 1979; Thorpe 1994; Devlin and Nagahama 2002). Consistent with the complex associations of growth with other traits is the finding that the genetic architecture of this trait is typically polygenic and determined by a complex network of genes (Filteau *et al.* 2013; Wellenreuther and Hansson 2016). This complexity makes it challenging to understand the specific genetic and physiological determinants that underpin faster growing phenotypes that need to be targeted when selectively breeding for enhanced growth.

For many commercially valuable species, selective breeding programmes have been initiated to produce strains that have an improved tolerance to domestic conditions and develop more quickly into a marketable phenotype (Olesen *et al.* 2003; Bernatchez *et al.* 2017). Understanding how domesticated organisms have been transformed from their ancestral wild type is valuable both from a genetic and evolutionary perspective, and provides fundamental information for future enhancement of strains through selective breeding. Genetic differences between wild and domesticated individuals can arise in three main ways. First, they can be inevitable by-product from a relaxation of natural selection pressures in captive conditions (Hutchings and Fraser 2008). By virtue of being raised in an artificial and controlled setting, farmed populations undergo inadvertent genetic changes because fish experience stable temperatures and no natural predators or significant foraging challenges. Second, intentional domestication selection (e.g. to enhance commercially relevant traits) and inadvertent or novel natural selection (e.g. the amount and quality of space provided) tend to favour fish that survive best in domesticated conditions, which can affect linked characteristics. Third, the often small population sizes of farmed species can cause strong genetic drift and inbreeding which can reduce the level of genetic diversity, increase the frequency of deleterious alleles, or overwhelm the strength of artificial selection and eliminate commercial important variation (Taberlet *et al.* 2011). This third process is commonly observed in the rapid loss of standing genetic variation in domesticated strains following a few generations of selective breeding (Hill *et al.* 2000).

Due to the recent domestication history of many fish species, they provide an ideal model for investigating the genetic changes associated with domestication and captive breeding because there are still natural populations that can be used as a reference (Mignon-Grasteau *et al.* 2005). Several studies to date have investigated the effects of domestication on fish gene expression patterns (Roberge *et al.* 2005; Devlin *et al.* 2009; Sauvage *et al.* 2010), but these are almost entirely focussed on salmonid fishes. Moreover, much less work has been conducted to dissect the more specific changes responsible for accelerated growth in selectively bred strains. Indeed, while the factors and pathways underlying differential growth in mammals have been established in considerable detail, knowledge of the relevant genes involved in growth variation in fish is much more limited (Fuentes *et al.* 2013). This is due to the relative lack of fish-specific molecular tools and functional studies, and compounded by the extra level of complexity in fish genomes due to whole genome duplication and the subsequent rediploidization event(s) (Volff 2005).

Here we explore temperature induced growth and stress responses in wild and domesticated strains of the Australasian snapper *Chrysophrys auratus* to gain a better understanding of the domestication process and identify the genes and pathways important for growth in teleost fish. This marine species has a distribution from 25 to 40° S in temperate and sub-tropical waters in New Zealand and Australia and is highly valued by commercial and recreational fisheries (Parsons *et al.* 2014). Despite its commercial and recreational importance, no transcriptomic studies have been performed on this species so far. We performed a manipulative experiment and held wild and domesticated snapper in cold and warm treatments and measured their growth responses and white muscle RNA expression profiles. First, we compare phenotypic responses of wild and domesticated snapper to the temperature treatments to quantify any growth related changes. Second, we compare gene expression profiles to identify the genes that are affected by genotype (wild *vs*. domesticated) and temperature, and their interaction. Finally, we use co-expression network models to gain insights into the metabolic modules and pathways affected in these genotype specific temperature responses, to better untangle growth and metabolic pathways in this species.

## MATERIALS AND METHODS

### Fish holding and experimental setup

The Institute of Plant and Food Research (PFR) in New Zealand has been breeding *C. auratus* since 2004 in Nelson, and in 2014, a selective breeding programme was started to select for enhanced growth in this species. Experiments were carried out on two-year-old wild-caught and hatchery reared domesticated *C. auratus*, from December 2015 to January 2016 at the PFR Seafood Facility in Nelson. Wild *C. auratus* were captured by trawl during a research voyage of the vessel FV Bacchante in the inner Tasman Bay (centred on S41 12.700 E173 09.400) in February 2015 and transferred to the PFR facility the same day (and thus had time to acclimatise in the hatchery for at least 10 months prior to the trial). The domesticated *C. auratus* strain was raised from naturally spawned larvae propagated from wild caught broodstock held at the PFR Nelson Seafood Research Facility. Prior to experimentation both populations were acclimatized in separate 5000 litre tanks provided with flow-through filtered seawater at ambient temperatures and light conditions. Both wild and domesticated *C. auratus* were fed on equivalent rations of mixed fish pieces, commercial aquaculture pellet (3 mm, Nova ME; Skretting, Cambridge, Tas., Australia) and an in-house formulated gel diet consisting of 21.3% Protein, 2.7% Lipid, 5.6% Carbohydrate, 7.7% Ash, 62.7% moisture. All rearing, holding and sampling procedures were performed using standard hatchery practices in accordance with New Zealand’s Animal Welfare Act (1999).

Twenty fish from each of the wild and domesticated strain (also referred to as genotypes) were weighed, measured and imaged, under anaesthesia (20ppm AQUI-S®, AQUIS NZ Ltd, Lower Hutt, New Zealand) and moved into four 800 litre tanks (2 × 10 wild-caught and 2 × hatchery reared *C. auratus*) supplied with 1μm filtered, UV sterilised, temperature-controlled flow through seawater (35ppt salinity, ambient temperature = 17.0°C). Once split, each tank consisted of either ten wild caught or domesticated *C. auratus* individuals.

Different thermal regimes were generated following a five day period of preconditioning at a nominal temperature of 17.0°C. One part of the wild caught and one domesticated strain were exposed to a temperature decrease of 1.0°C day^-1^ while the remaining other wild and domesticated strain were exposed to a temperature increase of 1.0°C day^-1^. Following the five day period of increased/decreased temperatures the desired temperature differential of either 13.0 or 21.0°C was established, henceforth referred to as low and high temperature treatments, respectively. These temperatures were chosen to reflect seasonal differences that these strains would experience in their local environment (i.e. winter and summer temperatures). Temperature loggers (HOBO, Onset Computer Corporation, MA, USA) showed that the thermal environment for the duration of the experiment maintained the desired treatment temperatures, with the low treatment having a mean temperature of 13.8°C and the high treatment having a mean temperature of 21.9°C, with minimal variation across the experiment (absolute maximum differences in the low and high treatment ±1.7°C and ±0.5°C, respectively).

Throughout the experiment, fish were maintained solely on the commercial pelletized (Skretting) diet described above at a ration equivalent to 2% body-mass per day, provided on a daily basis. Dissolved oxygen levels were checked multiple times per day to ensure levels were kept at >90% and tanks were cleaned every 3-4 days. Once the desired temperatures were reached, the experiment was allowed to run for 18 days.

### Animal sampling, RNA extraction and sequencing

Upon termination of the experiment, eight fish from each treatment were anaesthetised (25ppm AQUI-S^®^) then netted from their tank and subsequently euthanized with an overdose of anaesthetic (60ppm AQUI-S^®^). Immediately following euthanasia, fish were imaged and weighed for identification and phenotyping. Then a 2-3ml sample of mixed whole blood was collected from the caudal vein and placed on ice using 21g hypodermic needles and EDTA treated vacutainers (BD, Franklin Lakes, NJ, USA). For the transcriptome assembly, 12 brain and 12 epaxial white muscle tissues muscle (removed from the D-muscle block immediately anterior of the dorsal fin), as well as three whole larvae (2-3 months of age) were preserved in RNAlater (Ambion, USA) at 4°C overnight before being transferred to −80°C for long-term storage.

Haematological parameters were assessed in all individuals to compare the physiological conditions between the two treatments. Haematocrit (Hct) was measured from whole blood immediately after collection using 75mm capillary tubes spun at 12,000rpm (4°C, 5min, Heidelberg, Germany). Haemoglobin concentration [Hb] was then measured spectrophotometrically by mixing 10μl of whole blood into modified Drabkins reagent, measured in 1ml cuvettes (Wells *et al.* 2007). Mean Corpuscular haemoglobin concentration (MCHC) was estimated from the ratio of ([Hb]/Hct). Plasma was then separated by centrifugation (4,600rpm, 4°C, 10min) in 200uL aliquots, snap frozen in liquid nitrogen and then stored at −80°C for later analysis of plasma metabolites. Plasma osmolarity was determined from freeze-thawed plasma samples using a Wescor vapour pressure osmomter (Vapro 5520, Wesco Inc. UT, USA). Plasma triglycerides were determined on a clinical blood analyser (Reflotron, Roche, Germany) using standard methods. Plasma lactate and glucose were measured using commercially available enzymatic assays kits (Megazyme K-Late and K-Gluc, Food Tech Solutions, New Zealand) performed and analysed in a 96-well microplate format (Clariostar, BMG Labtech, Germany).

Total RNA was extracted from 12 fish (six fish from each treatment) using the Trizol LS Reagent (Life Technologies) according to manufacturer’s instructions. RNA samples were individually prepared (including mRNA enrichment with a ploy(A) method) for sequencing using the Illumina Tru-Seq kit on two lanes of an Illumina HiSeq 2000 sequencer (paired-end 100bp sequencing, 160bp insert length, see Supplementary Table 1) at the Beijing Genomics Institute Shenzhen, China.

### Sequence data processing and *de novo* transcriptome assembly

All samples were used for the transcriptome assembly, but only the white muscle samples from the temperature experiment were used for the gene expression study. Sequences were first quality trimmed (trailing: 20; lowest quality: 30) and minimum length (< 60 bp) using Trimmomatic v0.36 software (Bolger *et al.* 2014). Trimming also included removal of putative contaminants from the UniVec database (https://www.ncbi.nlm.nih.gov/tools/vecscreen/univec/). Trimmed sequences were further quality checked with FastQC v0.11.5 software (https://www.bioinformatics.babraham.ac.uk/projects/fastqc/). At this step, one individual was removed because of lower sequencing quality.

Paired-end reads were assembled into transcripts (min length 200 bp) using the Trinity v2.2.0 *de novo* assembly pipeline (Haas *et al.* 2013) with a default k-mer size of 25-bp. Raw transcripts (242,320) were filtered for presence of Open Reading Frames (ORFs) (length ≥ 300 nt), longest isoform matches and mapping rate (≥ 1 TPM), following (Pasquier *et al.* 2016) procedure. The remaining transcript sequences were searched against Uniprot-Swissprot database (blastX; e-value < 10e-6). For quality checks, the *de novo* transcriptome completeness was assessed with BUSCO v1.1b metazoa database (Simão *et al.* 2015). We used TransRate v1.0.3 quality statistics to validate each transcriptome filtering steps (Smith-Unna *et al.* 2016).

### Differential gene expression and genotype x environment interaction

We first investigated genotype × environment interaction (GEI) from gene expression levels using a GLM approach (temperature × genotype) with the *‘glmFit’* function and a likelihood-ratio test implemented in the R package edger (Robinson *et al.* 2010). We only considered genes with false discovery rate (FDR) < 0.01 to be significant. To further quantify the additive effect of temperature and genotype, we conducted a GLM approach (temperature + genotype) on the dataset with prior removal of gene with significant interaction term. Genes were considered significantly expressed when FDR < 0.01 and |logFC| ≥ 2 (e.g. a fourfold difference between treatments).

### Co-expression network analysis

Signed co-expression networks were built using the R package WGCNA following the protocol proposed by Langfelder and Horvath (Langfelder and Horvath 2008) based on normalized log-transformed expressions values. The main goal of this analysis was to cluster genes in modules associated with genotype and temperature effects and relevant gradients of clinical traits. Briefly, we fixed a soft threshold power of 22 using the scale-free topology criterion to reach a model fit (|R^2^|) of 0.81. The modules were defined using the *‘cutreeDynamić* function (minimum 30 genes by module and default cutting-height = 0.99) based on the topological overlap matrix and a module Eigengene distance threshold of 0.25 was used to merge highly similar modules. For each module we defined the module membership (kME, correlation between module Eigengene value and gene expression values). Only modules with an absolute Pearson’s correlation value (|R^2^|) > 0.75 with temperature and genotype factors and with p-value < 0.001 were conserved for downstream functional analysis. For module visualization we selected the top 30 genes (hereafter called hub genes) based on the kME values. The resulting gene networks were plotted with Cytoscape v3.5.1 (Shannon *et al.* 2003).

### Gene ontology and KEGG pathway visualization

Gene enrichment analysis were conducted using GOAtools v0.6.5 (Klopfenstein *et al.* 2018) based on the go-basic database (release 2017-04-14). Our background list included genes used for the gene network construction (after removing low-expressed genes, n = 13,282; see section above). Only GO terms with p-adj < 0.05 and including at least three genes were considered (Supplementary Table 2). We matched each DEG to corresponding KEGG pathway via the online web server KAAS (Moriya *et al.* 2007). The respective KEGG pathways were plotted using the Pathview R package (Luo and Brouwer 2013).

## RESULTS AND DISCUSSION

### Phenotypic changes

Upon termination of the experiments marked phenotypic differences were observed between genotypes and temperature treatments (Table 1, Figure 1). At the beginning of the trial we measured slight and non-significant differences in starting mass and length between the wild and domestic *C. auratus*, and at the end of the trial, these differences increased and were statistically significant, favouring the domestic genotype (Table 1). Temperature-related growth differences were also visible with either genotype achieving a significantly larger growth (length and mass) gains in the high temperate treatment (Table 1). Moreover, low temperature had a profound effect on growth rate, with a net effect of near-zero growth in the domestic strain and negative growth (a reduction in mass) in the wild. This may indicate that the domesticated strain is more resistant to cold stress than the wild strain, but also is more responsive to increases in temperature, each of which benefits maintenance and growth respectively.

**Table 1:**
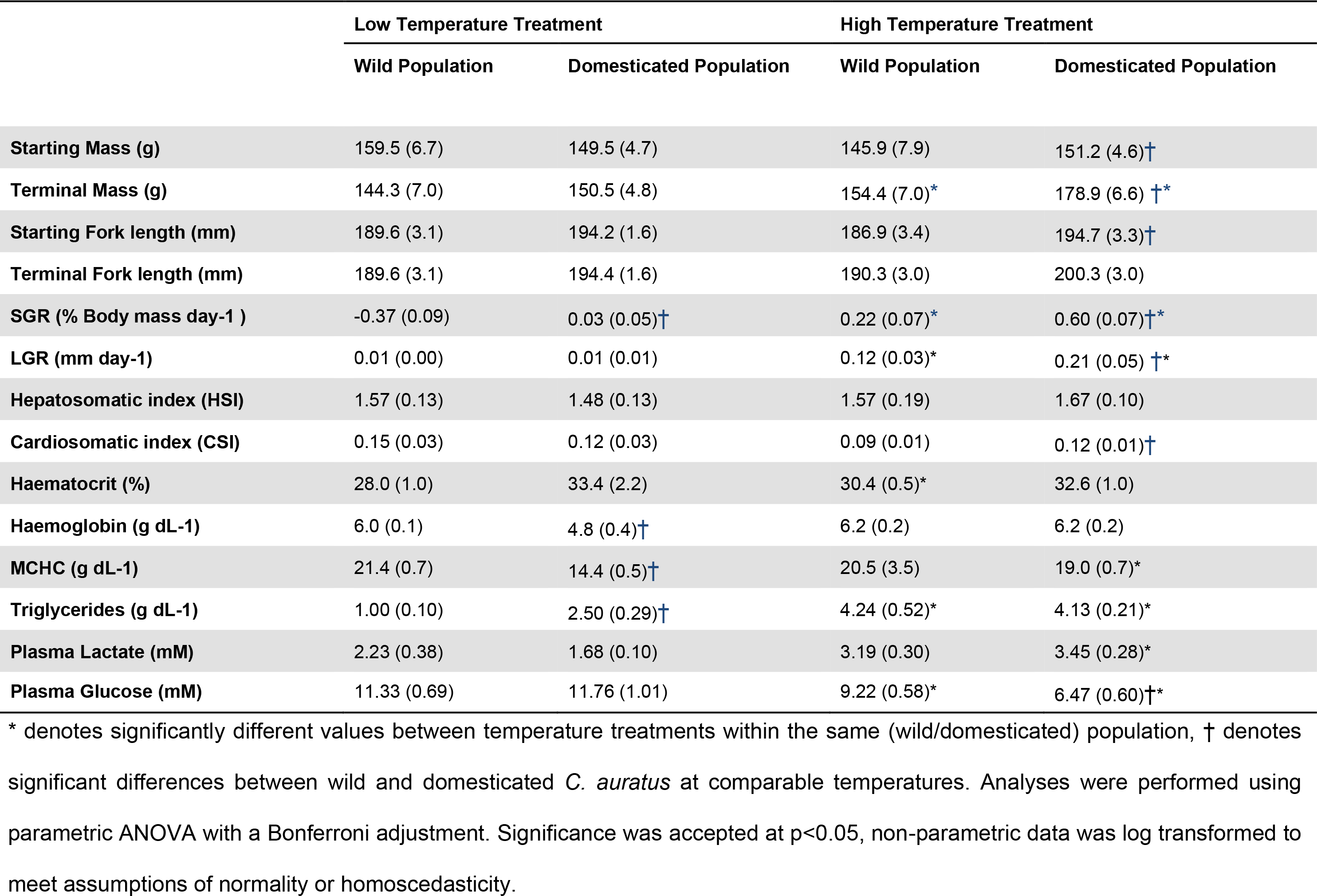
Phenotypic values of wild and domesticated *C. auratus* used in the low and high temperature treatments. Errors in brackets are sem, n=8. All physiological traits were measured upon termination of the experiment.

**Table 2:**
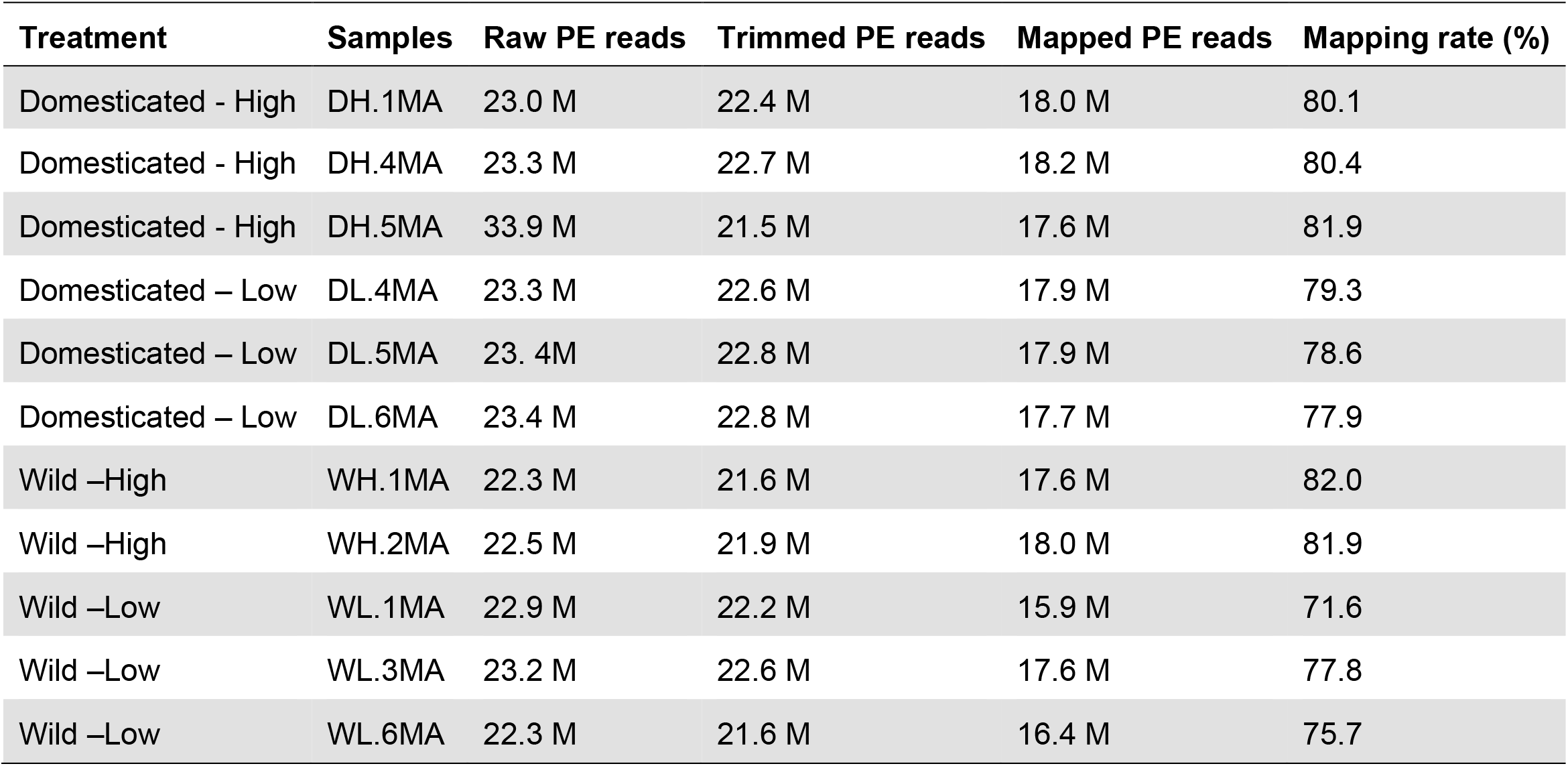
Sequencing results, reads pre-processing and mapping summaries.

**Figure 1:**
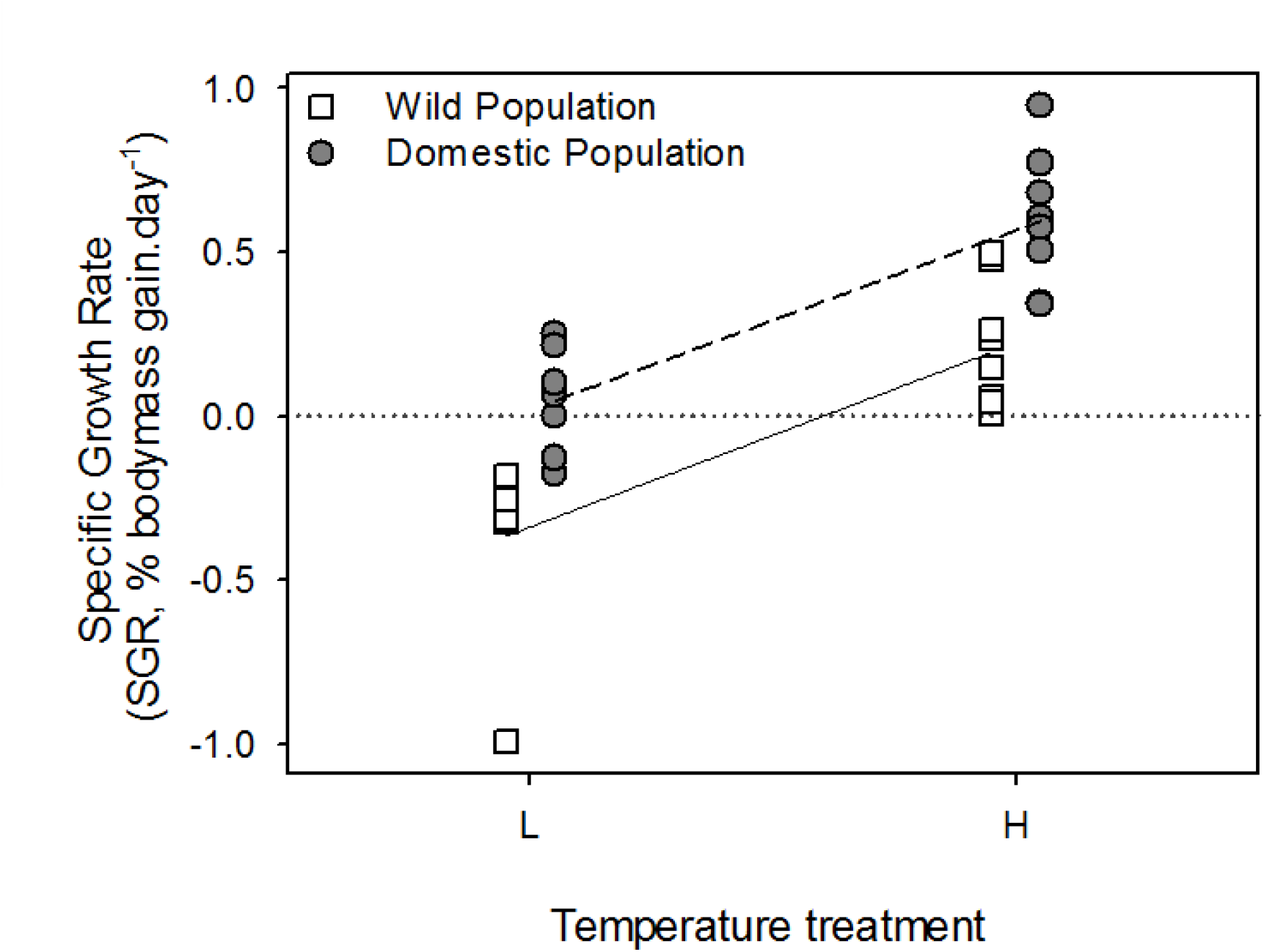
Specific growth rates of wild and domesticated *C. auratus* (8n/treatment) at the start and end of the low and warm temperature experiment.

### Transcriptome assembly quality and completeness

We used a total of 2.2 Gb paired-end reads to assemble a raw transcriptome containing 242,320 transcripts (215 million bases). The final reference assembly (after filtering) based on the different tissues types and replicate individuals represents a set of 33,017 transcripts (N50 = 2,804; GC content = 48.6%; Table 3). The final transcriptome completeness evaluation with BUSCO v1.1b (Simão *et al.* 2015) indicated that 96% of the highly-conserved single-copy metazoan genes (n=978) were present in the transcriptome sequence (including 89.5% complete and present a single-copy). Transcriptome annotation resulted in 26,589 transcripts matching 17,667 Uniprot-Swissprot entries (e-value < 10^-6^; Table 3) for a total of 11,985 transcripts associated with at least one of the Gene Ontology terms. For downstream expression analysis, only genes actually expressed in muscle (log CPM < 1 in at least two individuals) from the final assembly were retained, resulting in a total of 13,282 transcripts.

**Table 3:** Assembly and annotation statistics

| Transcriptome statistics |  |
| --- | --- |
| Total number contigs | 33, 017 |
| Percent GC | 48.61 |
| Contigs N50 (bp) | 2,804 |
| Total assembled bases | 63, 545, 739 |
| Median contig length (bp) | 1,482 |
| Average contig length (bp) | 1,924.64 |
| Annotation |  |
| Contig with Uniprot-sp match (e-value $10^{-6}$) | 26, 589 |
| Contig with GO identifier annotation | 25, 837 |

### Gene expression effects associated with domestication

We found that 206 genes were differentially regulated between wild and first-generation domesticated *C. auratus* (FDR < 0.01; |logFC| > 2), with 150 genes being up- and 56 down-regulated in wild individuals (Table 4). The overall percentage of domestication affected genes is thus 1.5 % following one generation in the hatchery, which may include the effects of human artificial selection, founder effects, genetic drift and inadvertent selection due to the new rearing environment or to the selection of traits correlated to the traits of interest. In addition to that, it should also be noted that the differences in the environment of domesticated and wild fish during the larval and juvenile phase, could have effected the long-term gene expression, and that some of these have persisted even though fish were acclimatised to hatchery conditions for over 10 months before the trial started. The gene ontology analysis revealed that the most enriched GO terms were involved in a strong global defense response (GO:0006952) and immune response (GO:0006955) (Supplementary Table 2), both of which were more highly expressed in the wild strain. Interestingly, among the most strongly differentially expressed genes, we found that two serum amyloid proteins (A-1 and A-3) were heavily downregulated in the F_1_-domestic *C. auratus* fish. Proteins or mRNAs of the serum amyloid A family are highly conserved have been identified in all vertebrates investigated to date and function as major acute phase proteins in the inflammatory response (Uhlar and Whitehead 1999). The co-expression network analysis identified a single module with highly significant genotype correlation (|R^2^| > 0.75; p-value < 0.001), namely the darkorange2 module (R^2^ = 0.88). This module contained 94 genes and showed no significant enrichment, but tendencies for increased glutathione metabolic processes (GO:0004364) and transferase activity, transferring alkyl or aryl (other than methyl) groups (GO:0016765). Furthermore, the most negatively correlated module with genotype was the antiquewhite2 module (n = 1,227; R^2^ = −0.69), which showed enrichment for adaptive and innate immune response functions (GO:0002250 and GO:0045087, respectively), defense responses (GO:0006952) and a positive regulation of the Mitogen-Activated Protein Kinase (MAPK) cascade (GO:0043410) and ELK3 coding gene (antiquewhite2 module).

**Table 4:**
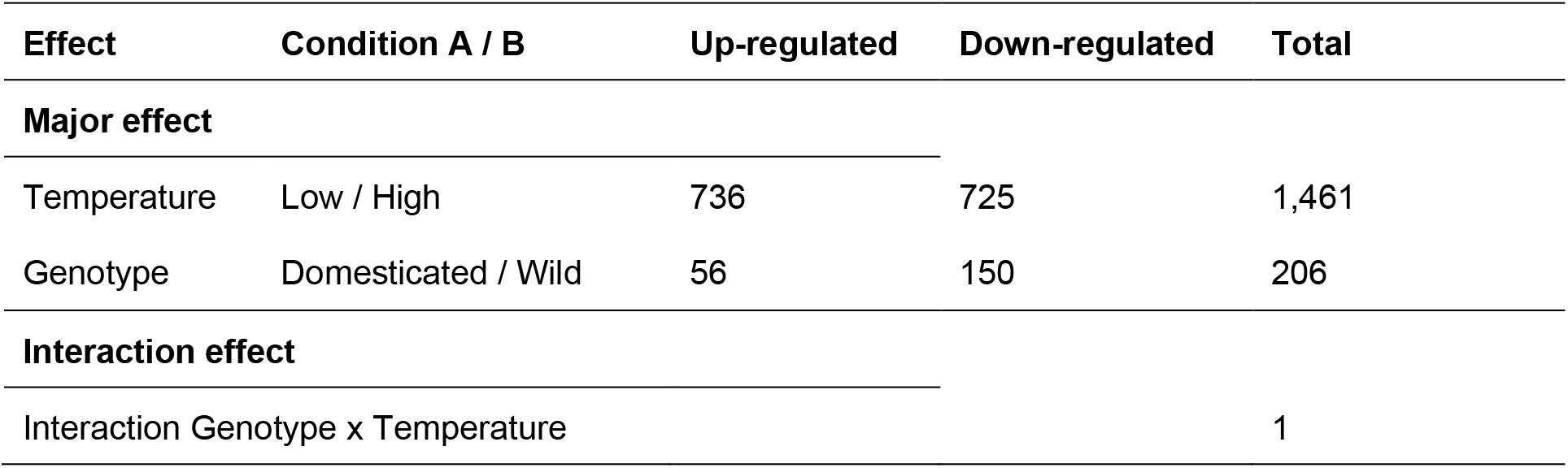
Number of differentially expressed genes using a contrast approach and glm model in edgeR. Genes were considered differentially expressed when FDR below 1% and |log2FC| > 2. A single gene showed significant interaction (genotype x temperature; FDR < 1%;).

| Effect | Condition A / B | Up-regulated | Down-regulated | Total |
| --- | --- | --- | --- | --- |
| **Major effect** |  |  |  |  |
| Temperature | Low / High | 736 | 725 | 1,461 |
| Genotype | Domesticated / Wild | 56 | 150 | 206 |
| **Interaction effect** |  |  |  |  |
| Interaction Genotype x Temperature |  |  |  | 1 |

In fish, the domestication process has been shown to influence metabolism, behaviour and chronic stress and immune response (Álvarez and Nicieza 2003; Millot *et al.* 2009; Douxfils *et al.* 2011a, 2015). In the first fish study on this topic, Roberge et al. (2006) showed that juvenile Atlantic salmon (*Salmon salar*) had gene expression differences for at least 1.4 and 1.7% of the expressed genes following 5-7 generations of domestication. Of the differentially expressed genes they found a general reduction in basal metabolic rate and an increased metabolic efficiency in farmed juvenile salmon compared to its wild counterpart, favouring allocation of resources towards growth and fat deposition. This finding is consistent with the faster growth and higher fat yield in domesticated salmon from the same source (Rye and Gjerde 1996). Interestingly, they also detected two genes coding for MHC antigens were more highly transcribed in some farmed salmon presumably in response to selection for disease resistance. In the perch (*Perca fluviatilis*) domestication increased the immune response during a challenge experiment with *Aeromonas hydrophila* with a congruent difference in the levels of HSP70 circulating between the first and fourth generation in captivity (Douxfils *et al.* 2011a, 2011b). With respect to somatic growth, domestication responses in Coho salmon (*Oncorhynchus kisutch*) closely resemble responses following growth hormone insertion seen in other fish species, most likely due to the strong selection for enhanced growth rate (Devlin *et al.* 2009). Indeed, recent studies on GH transgenic salmon showed that the growth advantage resulting from the GH insertion was tightly linked and dependent on the immune system response capacity, suggesting that the GH/IGF pathway interacts with the global immune response pathways (Alzaid *et al.* 2017).

Results from the current study were similar to the observations reported in previous studies, whereby differences in growth between domesticated and wild *C. auratus* occur through an interaction of the immune response and anabolic growth pathway modulation. Highlighted by the enrichment of immunity related processes and the increased expression of the MAPK/ERK cascade – a key regulator of IGF-I & II mediated myogenesis and somatic growth regulation in both mammals and teleosts (Codina *et al.* 2008; Fuentes *et al.* 2011) – these interactions suggest that in the wild *C. auratus* immune related activity was being prioritised above growth-related functions. This appeared to have negative consequences for the mass gain of the individuals over the experimental period. It is also noteworthy that modules segregating between genotypes are also tightly correlated to haematological indicators (Hb and MCHC outcomes, Table 1). When combined with the observation of green liver syndrome in the domestic low temperature treatment (results not presented), this outcome was considered to arise from a nutritional taurine deficiency in the experimental diets (Takagi *et al.* 2006; Matsunari *et al.* 2008), apparently exasperated by cold temperature exposure. It is interesting that this nutritional inadequacy interacts positively with the module enriched for immune response, and was not detected in wild individuals with the same recent nutritional history. This observation highlights the challenges of sourcing of nutritionally adequate feed for non-model as well as pre-commercial cultured fish species, as well as the unknowns implicitly associated with investigations involving wild caught fish. Additional studies will help to elucidate whether the patterns we observed result from different life history traits between genotypes (e.g. exercise, contact with pathogens, acclimation to captivity, different environmental and nutritional conditions in early life) and/or are the result of relaxed or novel selection in the early domestication processes.

### Temperature had a major effect on gene expression

To quantify the extent of gene expression variation associated with temperature and identify differentially expressed genes, we used a combined multivariate analysis (Redundant Discriminant Analysis; RDA) and GLM approach with an additive effect design (e.g. by removing the gene in the interaction, see Methods for details). Overall, temperature had the most profound effect on variation in expression explaining 47.2% of the total variance (Figure 2). This corroborates the differential expression analysis whereby (despite stringent filters FDR < 0.01; |logFC| > 2) we found a large number of genes (n = 1,461) differentially regulated between temperatures with 736 up- and 725 down-regulated genes in the high temperature relative to low temperature condition (Table 4; Figure 3). The gene ontology analysis on the total DE genes revealed that most enriched GO terms included tRNA aminoacylation for protein translation (GO:0006418) and sister chromatid segregation (GO:0000819), suggesting important differences in protein synthesis and cellular multiplication, both of which are key processes in myogenisis and somatic growth in fish.

**Figure 2:**
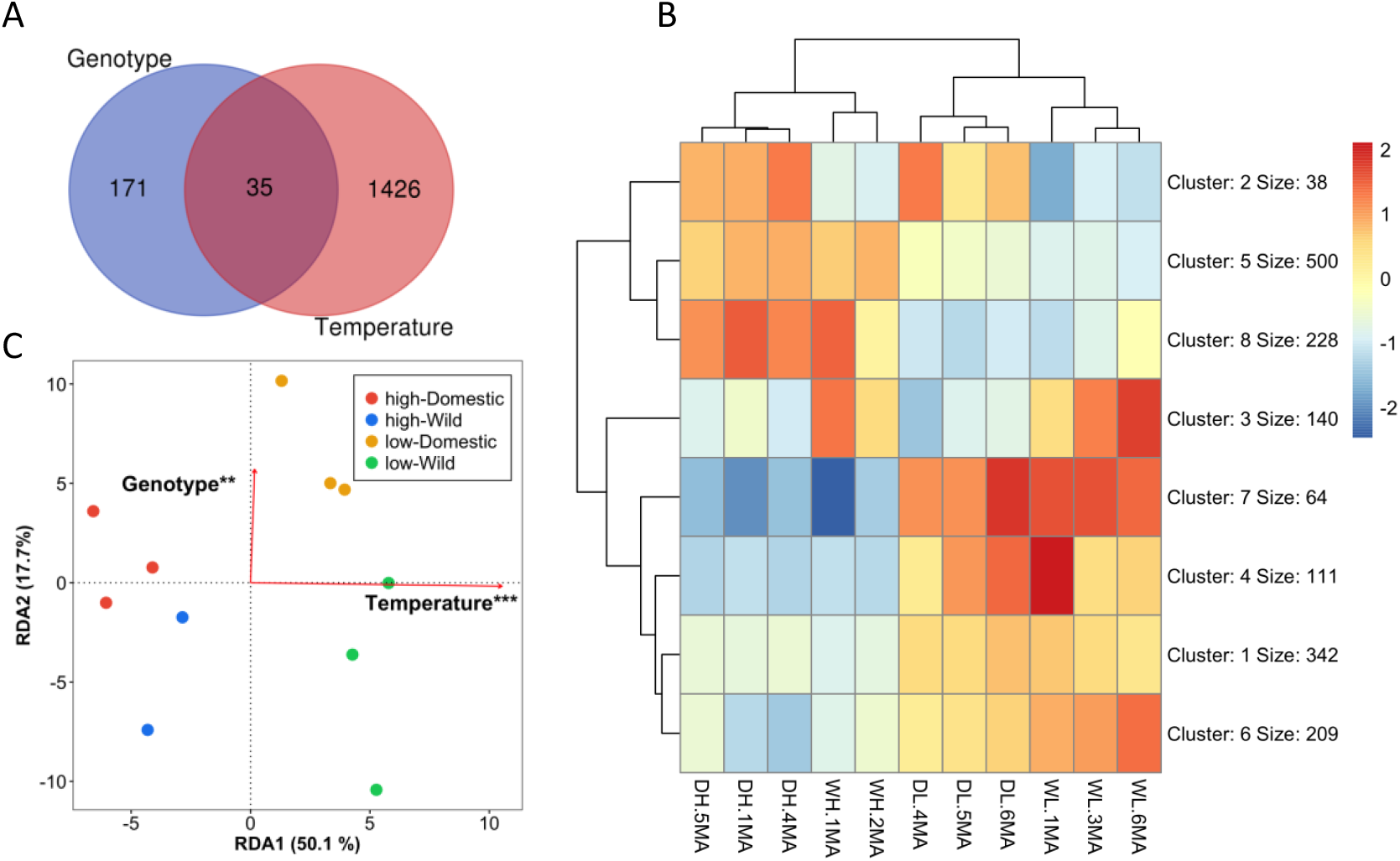
Effect of temperature and genotype one gene expression. A) Venn diagram showing the overlap between genotype and temperature in an additive effect (parallel reaction norms); B) Heatmap and K-means clustering of genes showing differential expression between genotypes and/or temperature; C) Distance-base redundancy analysis (db-RDA) performed on the expression data (logCPM [prior count 2)]. Only genes with min logCPM > 1 in at least 3 samples were retained for the analysis (n = 14,372). The db-RDA model was globally significant (p < 0.001) and explained 59.8% of all expression variation (adj. R^2^ = 0.598). Genotype and temperature significantly explained 17.2% and 47.2% of the variation, respectively, after controlling for each other with subsequent partial db-RDAs. Significance codes: p-value < 0.001 ‘***’; p-value < 0.01 ‘**’

**Figure 3:**
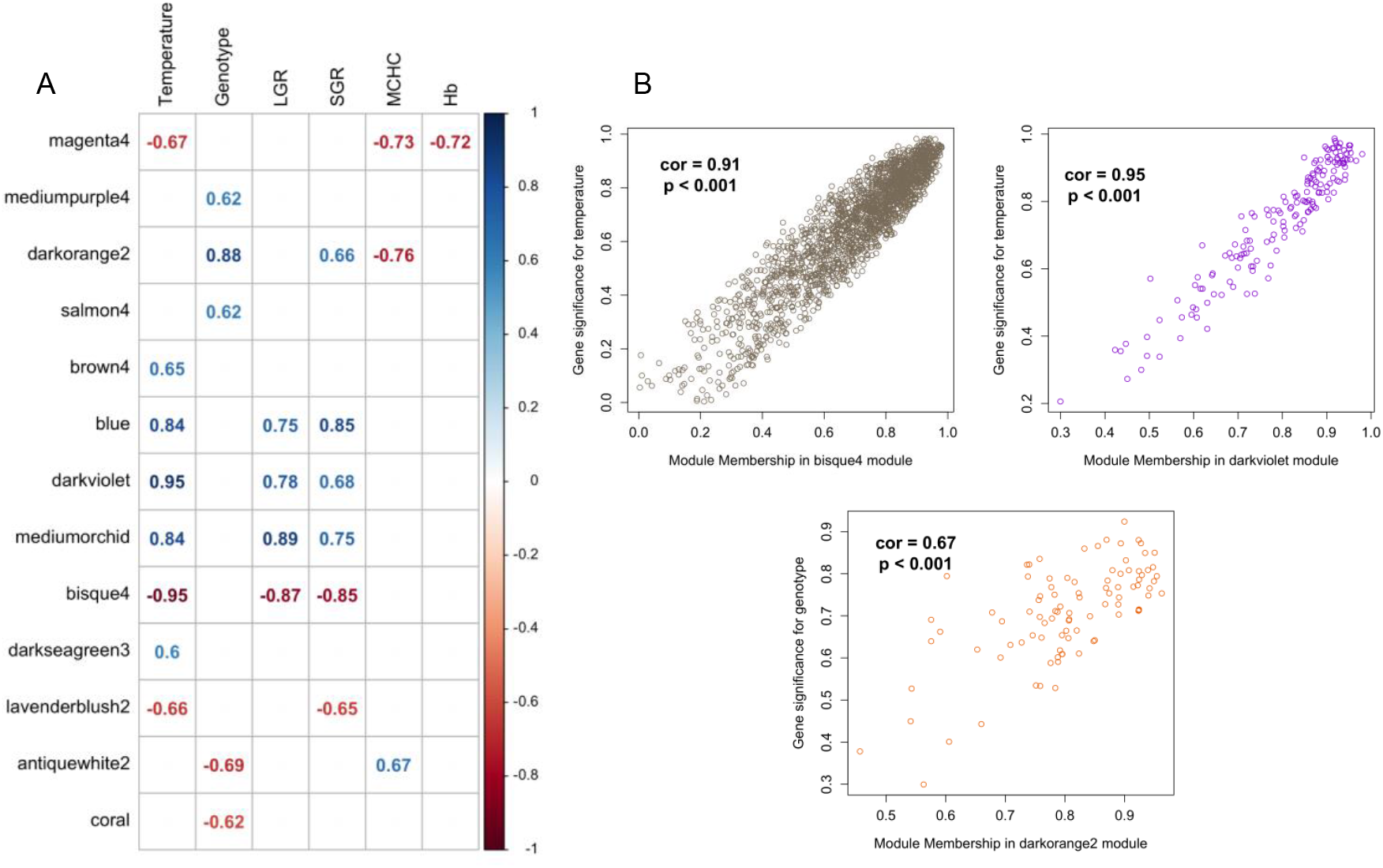
Co-expression network analysis. A) Correlation matrix from WGCNA. The matrix of co-expression was built on a total of 11,426 genes after removal of low-expressed genes (logCPM < 1 in at least 2 individuals and genes with gene expression variance < 0.1 in the global dataset). Only modules significantly correlating (p-value < 0.01) with temperature or genotype are represented. Values and indicate the correlation value (R^2^). B) Correlation between module membership and gene significance for the modules with the highest correlation to temperature (modules bisque4 and darkviolet) and genotype (module darkorange2).

We further dissected the global response to temperature using a co-expression network analysis, which has been shown to be particularly relevant for the functional analysis of non-model species (e.g. Filteau et al. 2013). We identified a total of 37 modules associated with a temperature response, but chose to only focused on 13 modules of these, based on whether they were significantly correlated (p < 0.01) with either temperature and/or genotype (Figure 3). A total of four modules showed a highly significant correlation (|R^2^| > 0.75; p-value < 0.001) with temperature, with three being positively [blue (0.84), darkviolet (0.95), mediumorchid (0.84)] and one being negatively correlated with temperature [bisque4 (−0.95)]. Four other modules, the magenta4, brown4, darkseagreen3 and lavenderblush2, showed a significant (p < 0.05) yet lower correlation with temperature, (R^2^ = −0.67, 0.65, 0.66 and −0.66, respectively). We validated the level of association by computing the mean gene significance value for each module and found that most correlated modules show the highest absolute mean gene significance to temperature (Figure 3; Supplementary Table 2). We also found that 81.4% of the DEGs between temperature treatments were for the four most strongly correlated modules in the WGCNA analysis, which was a finding consistent with our network construction. Finally, a gene ontology enrichment analysis was conducted for each module (gene ontology enrichment results are compiled in Supplementary Table 2).

We went on to investigate the role of genes in the co-expression network in the global transcriptomic response by identifying several major genes associated with rapid growth, thermal compensation and/or a post-acclimation response. The network co-expression analysis revealed that the four modules that were most responsive to temperature (blue, darkviolet, mediumorchid and bisque4) were also strongly correlated to SGR and LGR growth traits (Table 1), suggesting that these modules are either directly involved or linked to growth modulation, independently of the genotype. For the blue module (n = 1,404 genes) we again found significant enrichment (p-adj < 0.05) for amino acid activation (GO:0043038) and (GO:0043039) in accordance with our results for the DGE, but also for the generation of precursor metabolites and energy (GO:0006091), oxidoreduction coenzyme metabolism (GO:006733) and electron chain transport (GO:0022900). For the darkviolet module (n= 169 genes), we found significant enrichment for the haemoglobin complex (GO:0005833) and oxygen transport activity (GO:0005344). For the mediumorchid module (n = 1,370 genes), we found significant enrichment for global cell adhesion and metabolism, including extracellular structure organization (GO:0043062), collagen metabolic process (GO:0032963) and multicellular organism metabolic process (GO:0044236). Finally, the bisque4 module (n = 2,063 genes) showed enrichment for functions involved in peroxisome structure and activity including protein import into peroxisome matrix, docking (GO:0016560) and peroxisomal membrane (GO:0005778). We also found that the module brown4 (n = 93 genes) had enrichment for protein folding (GO:0006457), negative regulation of transcription from RNA polymerase II promoter in response to stress (GO:0097201), as well as tendency for a response to heat function (GO:0009408; p-value < 0.001; p-adj = 1). Gene ontology enrichment results are compiled in Supplementary Table 2.

We also identified hub genes, which are reported as core regulating genes, within the most relevant four modules correlated to temperature based on their modular membership (kME) values (Figure 3). The hub genes selection identified key actors in the global temperature response within each module, often known as key regulators on biological pathways. Among the blue module, we identified several t-RNA ligases but also the YTH domain-containing family protein 2 and the eukaryotic translation elongation factor 1 epsilon-1, that play a role in RNA protection following heat stress and DNA protection after damage, respectively (Lewis *et al.* 2017; Chen *et al.* 2018). Among the mediumorchid module we found cyclic AMP-dependent transcription factors (namely ATF-4 and 5) that are transcription factors associated with the circadian rythm regulation (Figure 4) as well as the Cryptochrome-1 coding gene, a core repressor of the circadian rythm (Hardie *et al.* 2012).

**Figure 4:**
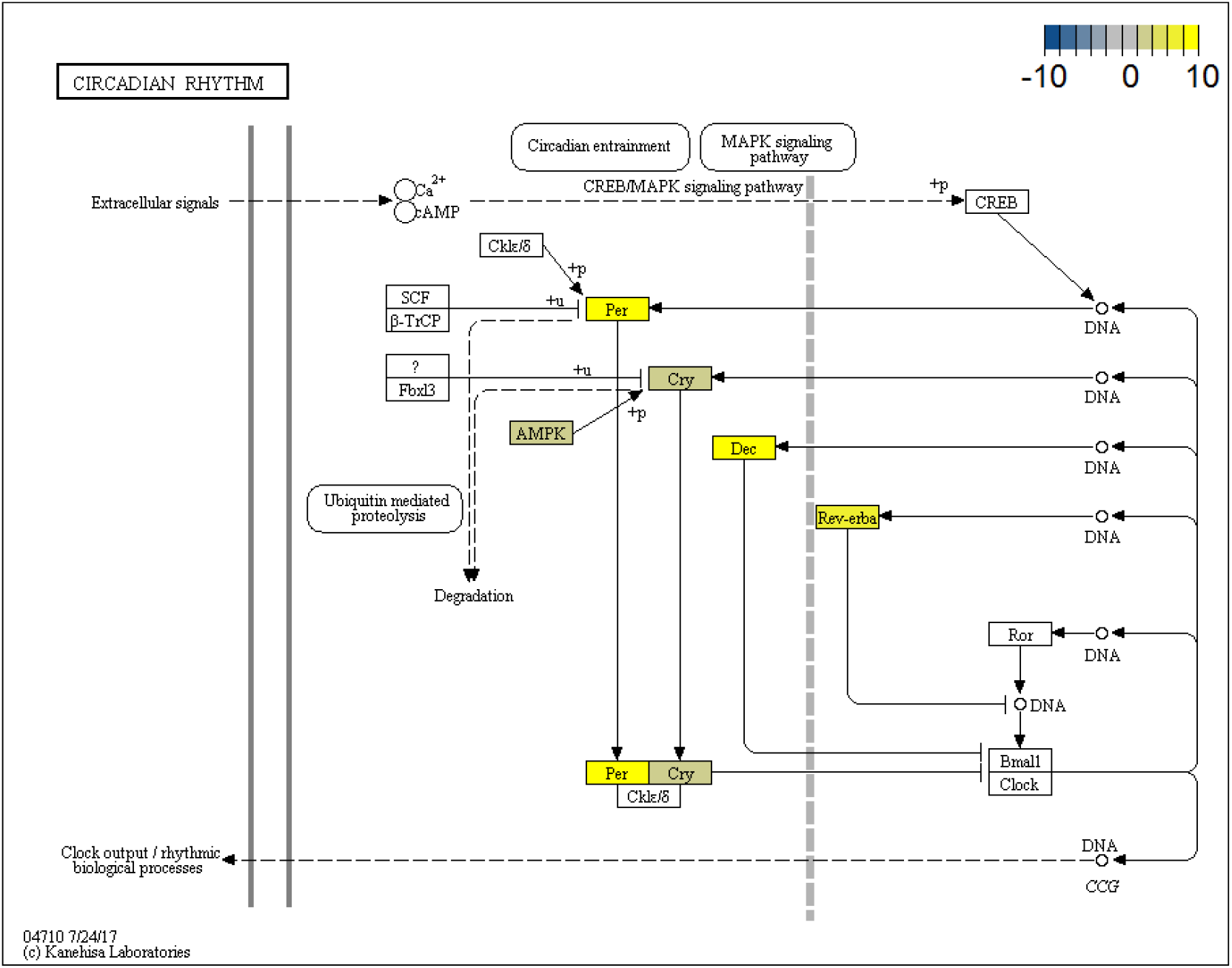
Differentially expressed genes in the KEGG pathway of the circadian rhythm. Kegg ontology were extracted from the KAAS online tools and plot using pathview package in R. Scale represent the log2FC expression levels between high and low conditions from edgeR analysis. Negative and positive values represent genes up-regulated and down-regulated in high temperature fish, irrespective of the genotype.

Given this extensive temperature-induced transcription level biological reorganisation, we focus the following interpretation on the most relevant results that corroborate previous transcriptional and physiological studies relating to the thermal responses of fish. HSPs-coding genes (HSPs90 and HSPs70 and HSP-binding70) were among the highest levels for differentially expressed genes we detected. Mostly clustered within the brown4 module (R2 = 0.65 with temperature), a significant increase in HSP expression was observed in *C. auratus* at temperatures at opposite ends of the species thermal envelope – at least in the geographic location the fish were cultured or captured in. This HSP response is well known in fish as a stress-induced response, which functions to protect against oxidative stress and apoptosis (Lindquist and Craig 1988; Iwama *et al.* 2004; Oksala *et al.* 2014). The HSP expression is often upregulated during short term (acute) exposures to high temperatures as seen in the gill and muscle tissues of the wild goby *Gillichthys mirabilis* exposed to 32°C over <8 hours (Buckley *et al.* 2006; Logan and Somero 2010). In addition, HSP upregulation can also be commonly observed during seasonally and environmentally relevant thermal regime shifts within the zone of sub-lethal thermal tolerance for a species (Fader *et al.* 1994; Oksala *et al.* 2014). The HSP response observed, clustered within the brown4 module, showed no significant correlation with growth (both SGR and LGR), nor were there detectable differences between wild and domestic strains. This is notable as differences in HSP expression commonly underlie phenotypic differences within a species, and this has often been observed across geographical gradients (Fangue *et al.* 2006; Hirayama *et al.* 2006).

Temperature produced a pronounced phenotypic effect characterised by a positive change to the growth rate at warm temperatures (~21°C), and negligible growth changes at low (~13C) temperatures. These phenotypic effects were underlined by substantial biological reorganisation associated with metabolic fuel switching and a shift from anabolic metabolism at high temperatures to maintenance/catabolism at low temperatures (Figure S3). Notable upregulation of AMPK (mediumpurple4 module) was evident in at low temperatures. Commonly referred to as the cellular ‘master switch’, this signaller is known to produce a cascade of changes to cellular homeostasis to reduce energetically expensive metabolic pathways (Hardie 2007). The increased expression of driver genes within the PI3K-AKT-mTOR pathway (darkviolet module) clearly underscore the up-regulation of cellular signalling pathways and growth-related processes (i.e. protein, lipid, and glycogen synthesis; cellular proliferation) at high temperature, all of which are known responses in fish (Fuentes *et al.* 2011, 2013). The upregulated PI3K-AKT activity together with the expression of FOXO1 (expressed within bisque4 module) corroborates the reported atrophic/catabolic processes that we observed during the low temperature conditions. Notably, these growth responses were also associated with increased expression of the Atrogin-1 muscle growth inhibitor and the downregulation of the muscle growth promoters IGFBP1&7 (also contained in bisque4 module) (Glass 2005; Fuentes *et al.* 2013). The observed switch from catabolic to anabolic states strongly suggests a switching of how the metabolic energy was being used in *C. auratus* at the two different temperatures. The most down-regulated gene at high temperature was the Long-chain fatty (LFA) acid transport protein 1, a gene involved in regulating LFA substrates in tissues undergoing high levels of beta-oxidation or triglycerides synthesis. The modulation of these transport processes corresponds with significant differences in circulating triglyceride levels (Table 1). Also associated with this process is the apparent ‘glucose sparing’ response and concomitant switch from carbohydrate to lipid based metabolism, inferred by the upregulation of both of beta-oxidation by ACC2 and gluconeogenic pathways via the PEPCK and phosphofructokinase / fructose bisphosphate mediated metabolic pathways (Figure S4). Similarly, down-regulation of N-terminal glutamine aminohydrolase - a hub gene of the mediumorchid pathway that favours the production of glutamate - was observed in the high temperature treatment. It is only that glutamate has recently been shown to present a major non-carbohydrate based energy substrate for skeletal muscle in fish (Weber *et al.* 2016; Jia *et al.* 2017).

The extensive reorganisation of *C. auratus* metabolism across their natural temperature range presents an interesting area of future research, particularly with consideration of the vast seasonally-dependent growth differences evident. Moreover, the molecular mechanisms underlying the notable thermal plasticity present in many fishes are still poorly understood. Differential expression of genes has been investigated as a cause for phenotypic plasticity in three-spine stickleback (*Gasterosteus aculeatus*) (Metzger and Schulte 2018). Being affected by both developmental temperature, as well as by adult acclimation temperature, there are probable mechanistic links between gene transcription, epigenetic signatures and, thermal plasticity across different time scales (Metzger and Schulte 2018). The plasticity in fish response to temperature may promote phenotypic alterations and ultimately, population divergence (Schulte 2014) during successive generations in both aquaculture and ecological contexts (Anttila *et al.* 2013; Donelson *et al.* 2018).

### Parallel and non-parallel reaction norms

A total of 35 genes showed parallel reaction norms whereby both wild and domesticated fish showed the same gene expression responses to temperature (i.e. significant effects of temperature and genotype; Figure 2). We further tested the hypothesis that temperature may impact gene expression but differently according to the genotype (interaction between temperature and genotype effects in a non-additive fashion; i.e. non-parallel reaction norms). To detect Genotype by Environment Interactions (GEI effects), we used a glm approach and a likelihood ratio test implemented in edgeR using the normalized data (Robinson *et al.* 2010). Only one gene, the Aryl hydrocarbon receptor nuclear translocator-like protein 2 (Baml2), showed a significant GEI (FDR < 0.01; Figure 5). The transcriptional activator Balm2 is a core component of the circadian clock regulation in mammals (Ikeda *et al.* 2000). Temperature-dependent activation and compensation of circadian rhythm have been observed in both vertebrates and invertebrates (Menaker and Wisner 1983; Sawyer *et al.* 1997; Rensing and Ruoff 2002; Zhdanova and Reebs 2006). Furthermore, different modulation of the circadian rhythm suggested adaptation to environmental cues after selective breeding for growth related traits during the early domestication process (López-Maury *et al.* 2008). Similarly, a switch in behaviour (day or nocturnal activity) has been observed during both temperature experiments and domestication selection in European sea bass (*Dicentrarchus labrax*) (Millot *et al.* 2009, 2010). The contrasting responses to temperature depending on the genotype background for some of the core regulators suggest that selection for a specific trait in aquaculture is also dependent of the rearing environment. More studies will be required to tease apart the responses to selection from any domestic environment-induced or plasticity effects that could occur during the larval rearing phase. Nevertheless, plasticity has significant impacts on the genetic gain calculation in many livestock breeding programs (Mulder 2016; Nguyen *et al.* 2017), and will be an important parameter to assess for newly domesticated species, including fish.

**Figure 5:**
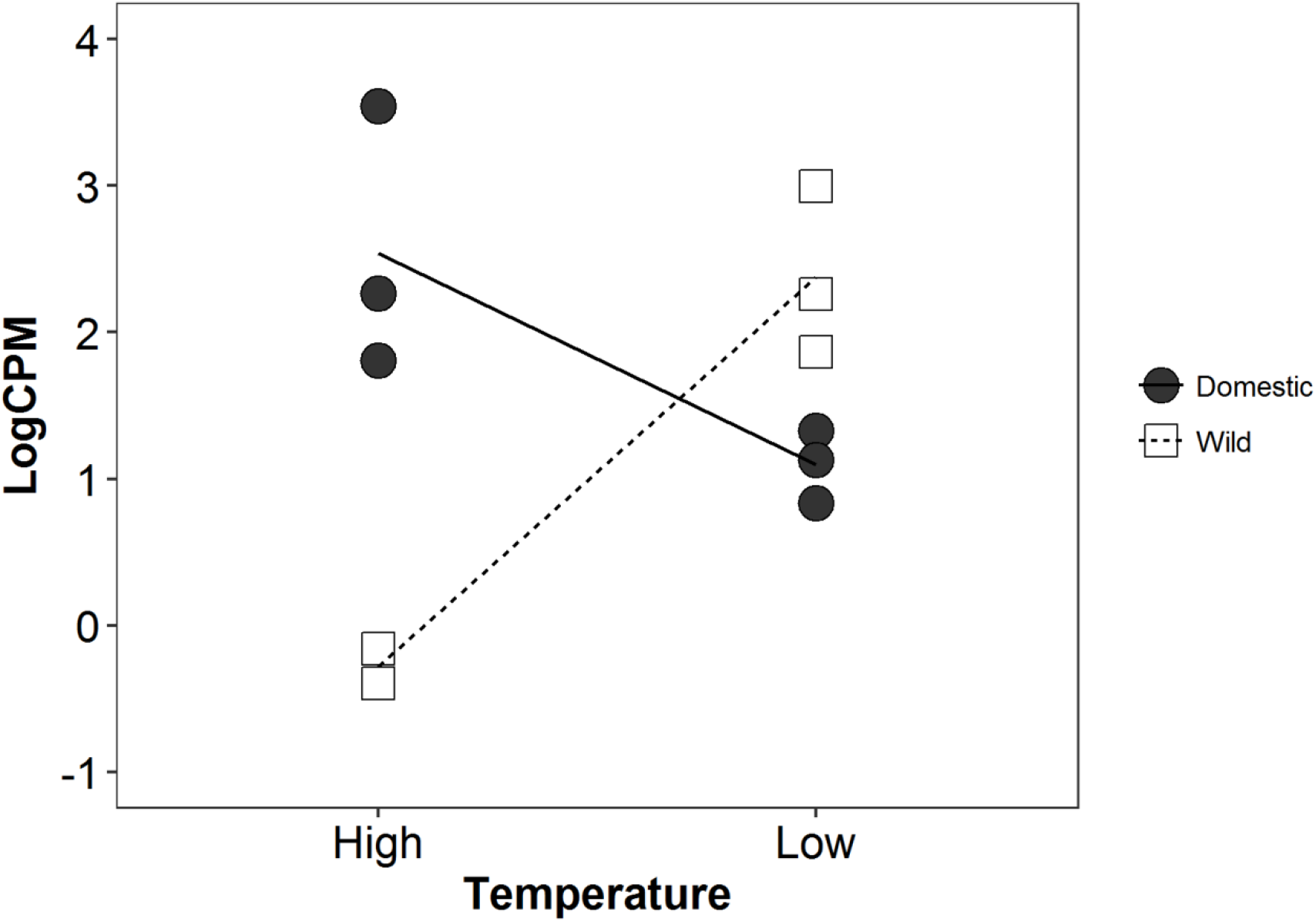
Gene interactions between temperature and genotype.

## CONCLUSIONS

Although fish domestication has received considerable interest for many years from a range of disciplines, modern large-scale genomic technologies and newly formed captive populations provide a unique opportunity to shed light on an important and well-documented evolutionary change for aquatic species. In this study, we combined GLM based approaches to investigate synergistic effects of recent domestication and temperature effects on gene expression regulation in the Australasian *C. auratus*. Coupling differential expression clustering and gene co-expression networks allowed us to begin to untangle the complex mechanisms of growth modulation during the first steps of selection and acclimation to domestication conditions. Our study shows that recent domestication and temperature had combined effects on muscle gene expression levels. We observed that temperature affected primarily HSPs responses as well as tissue development and cell turnover while genotype mainly affected the global immune response. Only a single gene (Baml2), crucial for circadian rhythm control, was affected by GEI. Admittedly, the present study only assayed a single tissue and further investigation at the brain or hormone producing tissue would produce a better understanding of the role of behavioural changes and immune responses that occur during the first few generations of domestication selection. Our study adds to the small number of previous studies that showed that gene expression responses can change rapidly following a few generations of domestication.

## DATA ACCESSIBILITY

The raw data were deposited on NCBI (*C. auratus* BioProject PRJNA484029) and will be accessible upon acceptance of the manuscript. For reproducibility, the codes are deposited in GitHub (https://github.com/jleluyer/PFR_snapper).

## AUTHORS’ CONTRIBUTIONS

MW and DC designed the experiments, MW and DC did the labwork, LB and JLL conducted the sequencing analysis. MW, JLL and DC wrote the manuscript. MW and PR developed the project proposals and funding. All authors read and approved the final manuscript.

## ACKNOWLEDGEMENTS

We would like to thank the staff at the Maitai Seafood Research Facility in Nelson, New Zealand, for help to run the experiment. This work was supported by a grant from the Ministry of Business, Innovation and Employment (MBIE) contract ID C11X1603 to MW and a VUW University Research Fund to PR. We would also like to thank Eric Normandeau for his help with genomic data analyses.

## SUPPLEMENTARY MATERIAL

**Table S1:** Sequencing, filtering and mapping statistics.

| Explanation | Read | length(bp) | Clean bases | Q20 (%) | GC (%) |
| --- | --- | --- | --- | --- | --- |
| Domesticated_High_1 | 46075138 | 46075138 | 99.37 | 99.1 | 49.66 |
| Domesticated_High_2 | 46697118 | 46697118 | 99.38 | 99.11 | 49.85 |
| Domesticated_High_3 | 44622780 | 44622780 | 98.87 | 99.12 | 50.73 |
| Domesticated_Low_1 | 46441522 | 46441522 | 99.39 | 99.09 | 50.24 |
| Domesticated_Low_2 | 46838422 | 46838422 | 99.38 | 99.1 | 50.02 |
| Domesticated_Low_3 | 46794464 | 46794464 | 99.4 | 99.13 | 49.69 |
| Wild_High_1 | 44578618 | 44578618 | 99.32 | 99.07 | 50.5 |
| Wild_High_2 | 45084660 | 45084660 | 99.4 | 99.11 | 50.22 |
| Wild_High_3 | 46410466 | 46410466 | 99.35 | 98.88 | 45.73 |
| Wild_Low_1 | 45726408 | 45726408 | 99.33 | 99.01 | 49.76 |
| Wild_Low_2 | 46485580 | 46485580 | 99.37 | 99.04 | 50.47 |
| Wild_Low_3 | 44561442 | 44561442 | 99.41 | 99.01 | 50.32 |

**Table S2:** Gene significance and module membership results for genotype and temperature and gene ontology results by module.

